# Simulating animal space use from fitted integrated Step-Selection Functions (iSSF)

**DOI:** 10.1101/2023.08.10.552754

**Authors:** J. Signer, J. Fieberg, B. Reineking, U. Schlägel, B. Smith, N. Balkenhol, T. Avgar

## Abstract

1. A standing challenge in the study of animal movement ecology is the capacity to predict where and when an individual animal might occur on the landscape, the so-called, Utilization Distribution (UD). Under certain assumptions, the steady-state UD can be predicted from a fitted exponential habitat selection function. However, these assumptions are rarely met. Furthermore, there are many applications that require the estimation of transient dynamics rather than steady-state UDs (e.g. when modeling migration or dispersal). Thus, there is a clear need for computational tools capable of predicting UDs based on observed animal movement data.
2. Integrated Step-Selection Analyses (iSSAs) are widely used to study habitat selection and movement of wild animals, and result in a fully parametrized individual-based model of animal movement, which we refer to as an integrated Step Selection Function (iSSF). An iSSF can be used to generate stochastic animal paths based on random draws from a series of Markovian redistribution kernels, each consisting of a selection-free, but possibly habitat-influenced, movement kernel and a movement-free selection function. The UD can be approximated by a sufficiently large set of such stochastic paths.
3. Here, we present a set of functions in R to facilitate the simulation of animal space use from fitted iSSFs. Our goal is to provide a general purpose simulator that is easy to use and is part of an existing workflow for iSSAs (within the **amt** R package).
4. We demonstrate through a series of applications how the simulator can be used to address a variety of questions in applied movement ecology. By providing functions in **amt** and coded examples, we hope to encourage ecologists using iSSFs to explore their predictions and model goodness-of-fit using simulations, and to further explore mechanistic approaches to modeling landscape connectivity.

## Introduction

Integrated step selection analysis (iSSA; Avgar et al. 2016; Fieberg et al. 2021) has emerged as a powerful and unifying methodological framework for quantifying different aspects of animal space use, including habitat selection patterns, movement behavior, and transient and steady-state utilization distributions (UDs). An iSSA results in a fully parametrized individual-based movement model that can be broadly classified as a locally-biased correlated random walk (Duchesne, Fortin, and Rivest 2015) which we refer to as an integrated step selection function (iSSF). In an iSSF, movement emerges from the product of a movement-free habitat selection function (MF-HSF; i.e., how would the animal select habitat if it were not constrained by movement) and a selection-free movement kernel (SF-MK; i.e., how would the animal move if it were not constrained by habitat selection). Note that the latter may include various habitat or environmental effects on movement, just not selection per se. Conceptually, the iSSF can be thought of as estimating a two-dimensional probability density function for the animal’s position after the next step (a redistribution kernel), given the environmental conditions and the animal’s current position and recent path. It provides a mechanistic model that can be fitted to data and used to simulate emerging patterns, the most basic of which is the UD (e.g., Signer, Fieberg, and Avgar 2017; Hofmann et al. 2023; Potts and Börger 2023).

Simulations from iSSFs have been used to investigate emergent patterns of space use from fitted iSSFs (Signer, Fieberg, and Avgar 2017; Potts and Schlägel 2020), to model connectivity between different animal populations or habitat patches (Hofmann et al. 2023; Whittington et al. 2022), and to evaluate and validate fitted models (Sells et al. 2023; Potts et al. 2022). Although analytical approximations of various estimation targets exist for some situations (Potts and Schlägel 2020; Potts and Börger 2023), simulations are more intuitive, flexible, and applicable to a wider range of problems. Despite the already widespread use and interest in simulations in movement ecology (Zurell et al. 2010; Whittington et al. 2022; Aiello et al. 2023), a general simulation routine is missing from available software, requiring analysts to write custom code. We address this gap by providing a user-friendly tool that can be used to address the various use cases described above.

We implemented two main functions, `redistribution_kernel()` and `simulate_path()`, in the **amt** package (Signer, Fieberg, and Avgar 2019) for the programming language **R** (R Core Team 2023). The first function computes a dynamic redistribution kernel from a fitted iSSF given a set of initial conditions (i.e., previous and current positions in geographic and environmental space). The second function is used to simulate movement paths by iteratively sampling a new position from a redistribution kernel and then updating this kernel to reflect the individual’s new position. We illustrate how simulations can be used to visualize different redistribution kernels, to generate data for various testing purposes, and to validate models and compute derived quantities (e.g., space use maps) in a case study using tracking data from an African buffalo. Finally, we discuss other applications that may be of interest to a wide range of ecologists.

## Methods

### Background

The iSSF can be used to calculate a redistribution kernel that gives the probability of moving to position *s* at time *t* + *τ* (*τ* being a constant time step), given the animal is at position *s*^*′*^ at time *t* and was at position *s*^*′′*^ at time *t* − *τ* . More formally, the value of the redistribution kernel *u*(*·*) for a tentative position s at time *t* + *τ* is given by

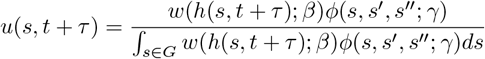

where *w*(*·*) is the MF-HSF and *ϕ*(*·*) is the SF-MK. The denominator normalizes the redistribution kernel over all possible positions *s* within the spatial domain *G*. The selection parameters, *β*, weigh different habitat attributes (sometimes referred to as ‘resources’), *h*, at position *s* and time *t* + *τ*, and the movement parameters, *γ*, are used to model the distribution of step lengths and turn angles.

When step lengths and turn angles are modeled using distributions from the exponential family, and an exponential MF-HSF is used, the numerator can be rewritten in log-linear form as

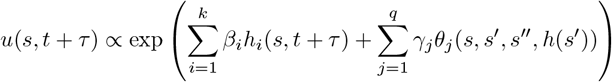

where *h*_*i*_(*s, t* + *τ*) is the value of the *i*-th (out of *k*) habitat attribute, *h*_*i*_, at position *s* and time *t* + *τ*, and *θ*_*j*_ the *j*-th (out of *q*) geometrical attribute of the step (e.g., the cosine of the turn angle, the step lengths, or the log of the step length, which are movement characteristics that depend on the assumed step-length and turn-angle distributions). The *θ*_*j*_ can also consist of, e.g., the product of the step length and the value of a certain habitat attribute at *s*^*′*^ to model environmental effects on movement. The parameters of the model can be estimated using different approaches. The most common method is a two-step procedure, estimating first tentative parameters for the SF-MK and using these to estimate the *β*_*i*_ while simultaneously updating parameters of the SF-MK (see Avgar et al. 2016; Fieberg et al. 2021, in particular Appendix C).

### Implementation

The **amt**-function `redistribution_kernel()` creates a redistribution kernel from the object returned by `fit_issf()`, using the two-step procedure. In situations where the parameters have been estimated in some other way (e.g., using Poisson regression Muff, Signer, and Fieberg 2020; or a full likelihood approach Schlägel and Lewis 2016), or when simulating from scratch based on user-defined parameter values, the necessary object can be created with the `make_issf_model()` function. The `redistribution_kernel()` function requires additional arguments, especially: `map`, `fun`, and `landscape` (Table 1). The argument `map` must be a `SpatRast` from the `terra` package (Hijmans 2023) and must contain all environmental covariates included in the model; its extent determines the extent of the simulation landscape. The argument `fun` is a function that is executed at each time step of the simulation to extract (and possibly manipulate) the values from map. Often, the default function – `extract_covariates()` – is sufficient. Finally, the argument `landscape` controls whether the redistribution kernel is implemented in continuous space and approximated using Monte Carlo sampling (`landscape = “continuous”`) or in discrete space (`landscape = “discrete”`). Generally, a stochastic redistribution kernel in continuous space is preferable; a discrete-space approximation can lead to a biased step length distribution, since the smallest step length is then given by the resolution of the environmental covariate raster. Continuous redistribution kernel can use the tentative step length and turning angle distributions as proposal distributions for stochastic simulations from the selection-free movement kernel. For visualization purposes, however, we may be interested in a discrete approximation of the redistribution kernel. In this case, we need to: 1) update the tentative step length and turning angle distributions to the SF-MK using coefficients associated with movement characteristics; and 2) account for the transformation from polar to Euclidean coordinates (see Supplement 1), the function `redistribution_kernel()` takes care of this.

Once multiple paths are simulated (each a stochastic realization of the same iSSF), they can be used to approximate either a transient utilization distribution or a steady-state utilization distribution (Signer, Fieberg, and Avgar 2017). A transient UD is a probability surface of animal occurrence at the end of all possible paths starting from a given point in space and time, and lasting a given duration. A transient UD is thus spatially and temporally specific – it takes different forms depending on the starting conditions and the sampling duration. For a given starting position (in space and time) and duration (= number of steps), the transient UD is approximated as the intensity of the point pattern formed by the endpoints of many simulated paths (the more simulated paths, the better the approximation). A steady-state UD is the probability surface of animal occurrence at the limit of an infinitely long path – it is independent of the initial conditions. A steady-state UD is thus approximated by simulating paths so long that the resulting point pattern of step endpoints is no longer sensitive to the starting point. Note that, since a steady-state UD is independent of duration, all simulated step endpoints are included in the summary (rather than just the last endpoint of each simulated path as in the transient UD). In cases where a single path cannot be expected to effectively visit all locations within the spatial domain in a computationally feasible time frame, multiple (long) paths should be simulated, each starting from a different starting point across the domain. Both types of UDs could be further smoothed using a kernel density estimator applied to the resulting point pattern (Potts and Börger 2023).

## Case Study

### Simulating movement from scratch

First, we show how our simulator can be used to visualize different redistribution kernels and simulate from them. For our model of step lengths, we used a gamma distribution with parameter values for shape = scale = 2. To model turning angles, we used a von Mises distributions with two different concentration parameters, 0 (no directional persistence) or 4 (strong positive directional persistence). Lastly, we allowed the individual to select for or against a spatially varying habitat attribute (gray square in Fig. 1). The overall redistribution kernel *u*(*s, t* + *τ*) for any steps starting at *s*^*′*^ and ending at *s* can be described as

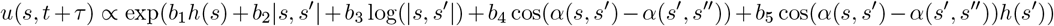

where *b*_1_ is a selection coefficient of the MF-HSF; *b*_2_, *b*_3_, *b*_4_ and *b*_5_ are parameters in the SF-MK, |*s, s*^*′*^| is the Euclidian distance from *s*^*′*^ to *s* (step length), *α*(*s, s*^*′*^) is the angular heading from *s* to *s*^*′*^, (*α*(*s, s*^*′*^) − *α*(*s*^*′*^, *s*^*′′*^)) is thus the turning angle relative to the previous step, and *h*(*s*) is the habitat value at a given position *s*. First, we generated six different redistribution kernels on a discrete landscape by varying the values of *b*_4_ and *b*_5_ to illustrate different SF-MKs (Fig. 1a, b, e, f) and *b*_1_ to illustrate different MF-HSFs (Fig. 1c, d).

**Figure 1:**
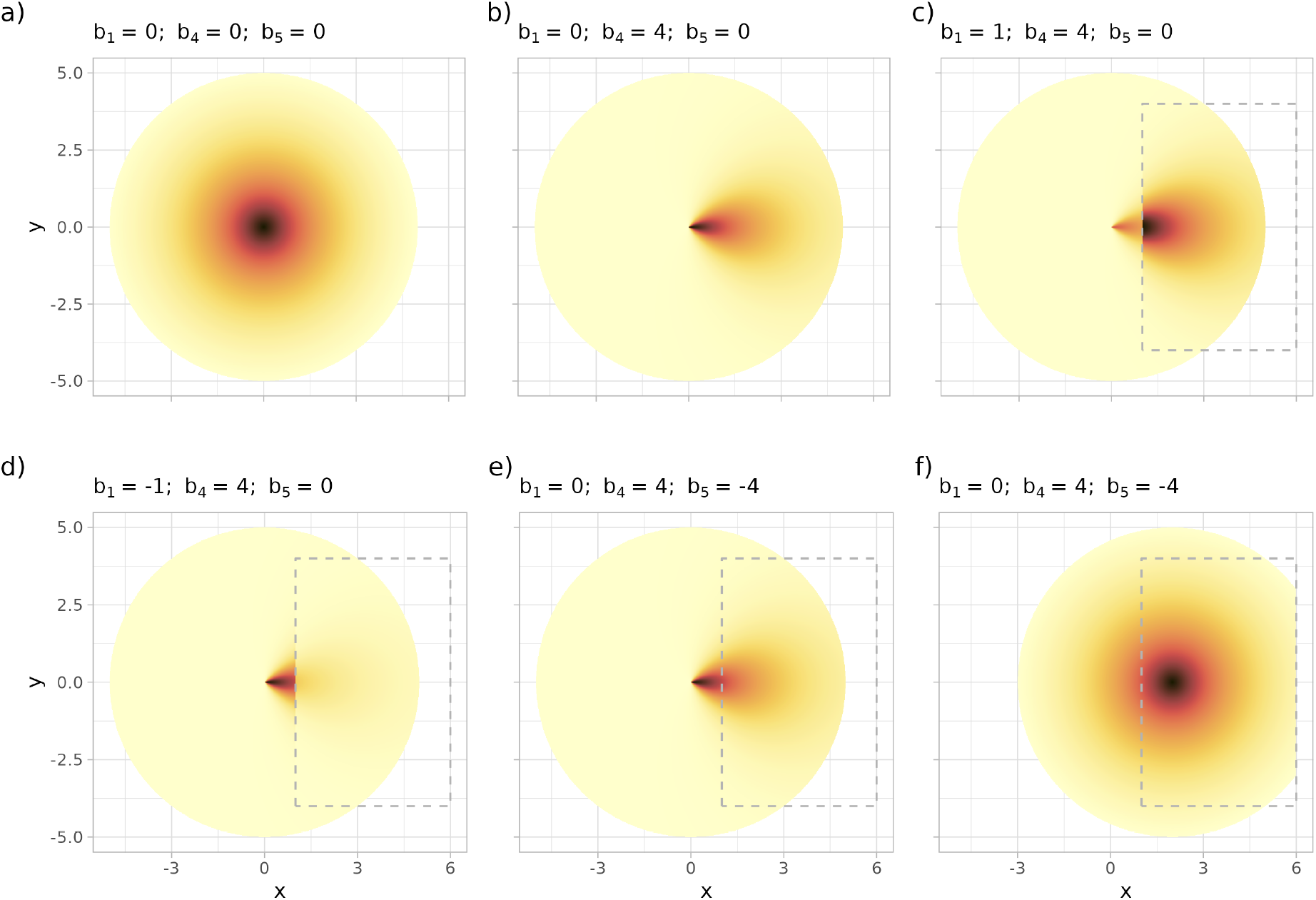
Different redistribution kernels resulting from different parametrizations of the selection-free movement kernel (SF-MK) and the movement-free habitat selection function (MF-HSF). In the simplest case, there is no habitat selection and movement is only constrained by the SF-MK, which excludes (panel a) or includes (panel b) directional persistence. An environmental covariate (gray rectangle within which h = 1, as opposed to out of the rectangle where h = 0) can lead to preference (panel c) or avoidance (panel d). Furthermore, the SF-MK can also depend on the habitat the animal is in at the start of the movement step. We show redistribution kernels for a case where the animal exhibits different directional persistence depending on whether it is located outside (panel e) or inside (panel f) the gray rectangle.

Second, we simulated 50 paths for 30 time steps each by repeatedly sampling from successive redistribution kernels (Fig. 2a). We assumed that the animal had little directional persistence and selected for habitat within the gray dashed rectangle (Fig. 2b). We then applied a kernel density estimator to the endpoints of these 50 paths (Fig. 2c) to obtain a smooth estimate of the transient UD (Potts and Börger 2023).

**Figure 2:**
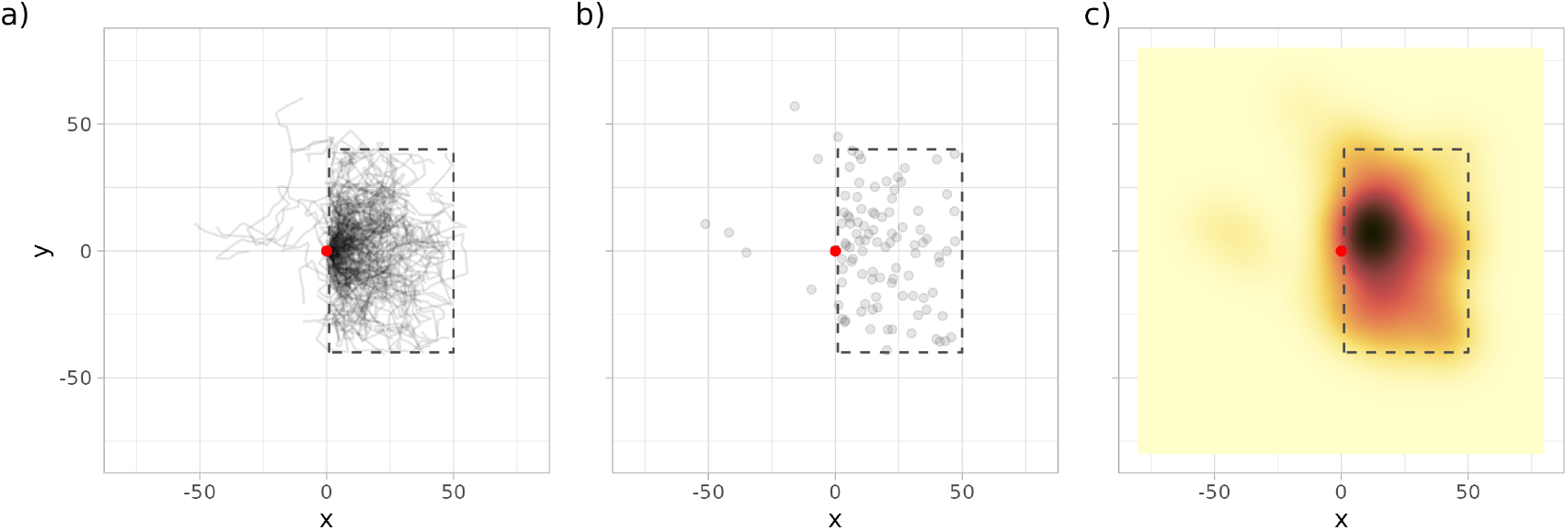
Figure 2: Simulated movement paths of 50 animals for 30 time steps (panel a). We then used the end positions (panel b) to generate a smoothed map representing the transient Utilization Distribution at *t* = 30 (panel c). The start point is marked with a red dot.

### African buffalo

We used tracking data from Cilla, an African buffalo, previously used to introduce the local-convex-hull home range estimator and freely available from Movebank (Getz et al. 2007; Cross et al. 2016). We fitted three iSSF models of increasing complexity. In the first, we modeled habitat selection as a function of distance to the nearest river at a spatial resolution of 90 m. Next, to model home ranging behavior, we added the x and y coordinates of the endpoint of each step (observed and control) and the sum of their squares (see Appendix S3 of Alston et al. 2023). Finally, we included the river as a potential barrier to movement. For each step (observed and control) we compared whether or not the start and end of a step were on the same side of the river. Data and reproducible code for all three models are provided in Supplement 2.

The African buffalo case study illustrates how simulations can be used to visually check model fit (Fig. 3). In model 1, movement is unconstrained and the animal frequently leaves the landscape (Fig. 3; left panel). In model 2, the inclusion of home ranging behavior constrains the animal to never leave the landscape, but it does not prevent the animal from crossing the river even though river crossings were never observed in the data (Fig. 3; middle panel). In model 3, the parameterized iSSF produces a much more realistic movement path (Fig. 3; right panel). Note that there are still unexplained patterns in the observed path (e.g., elevation could also be important), but we conclude that model 3 is already a significant improvement over model 1. Many realizations of this simulation could be used to formally measure the predictive power of each model (Potts et al. 2022).

**Figure 3:**
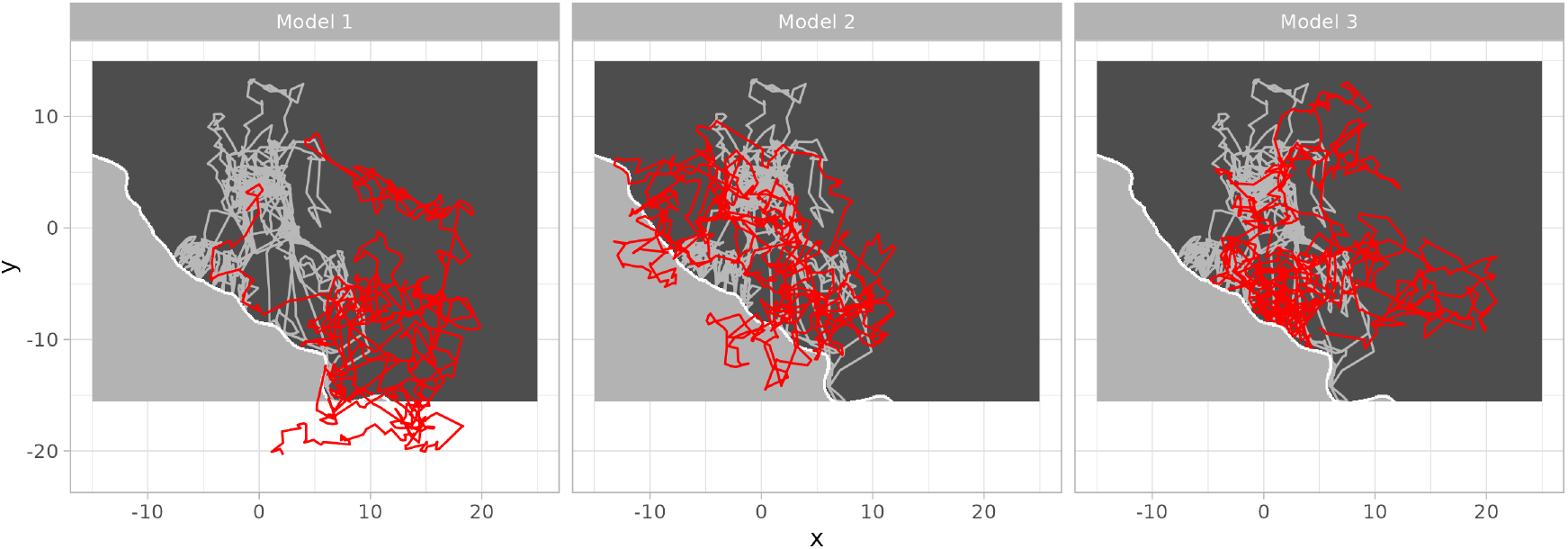
Figure 3 : Observed (light gray) and simulated (red) movement paths of an African buffalo. All models included distance to water as a covariate. Model 2 and Model 3 also included the coordinates at the end of each step to account for home ranging behavior. Model 3 also included whether the start and end points of each step were on the same side of the river (shown in white).

## Discussion

We have developed functions in R that enable users to simulate animal space use directly from fitted iSSFs using redistribution kernels that are dynamic in space in time. Our approach builds on an established workflow for data analysis.

We see several different applications for such a simulator, including model evaluation, prediction, and estimation of space use metrics (such as landscape connectivity) to inform conservation. First, simulated and observed paths can be visually compared (Fig. 3). If the model has been specified in a way that describes the data-generating process reasonably well, the observed path should not stand out among the simulated path. Similar to our case study, one can evaluate whether the observed and simulated paths exhibit similar behavior near roads, rivers, or other prominent environmental features. Second, our simulator can be used as a way to develop a null distribution to test for evidence of site fidelity and/or memory (Picardi et al. 2023). Third, predicting the steady-state or transient UD of an animal is often of interest. When the redistribution kernel is static (i.e., does not change spatially), other approaches are available to generate steady-state UDs (Signer, Fieberg, and Avgar 2017; Potts and Börger 2023). However, if the goal is to predict short-term, transient utilization distributions or if there is no steady-state UD (e.g., if the redistribution kernel is periodic in time), the simulator presented here offers a natural way forward. Finally, animal movement is of interest for many conservation applications and questions that require quantification of landscape connectivity. Unlike many current approaches, our simulator provides a way to explore connectivity via a mechanistic model of animal movement.

We have described the simulator in the context of simulating from a fitted iSSF, but it is also possible to simulate paths from scratch (as we did in the first case study). This requires the analyst to define step length and turning angle distributions for the movement model and selection coefficients for the selection functions. This feature makes our approach useful for exploring research questions via simulation or for evaluating different sampling designs.

We expect the recent interest in simulations from integrated step selection functions to continue. Recent extensions to iSSAs that include memory (Rheault et al. 2021), behavioral states (Klappstein, Thomas, and Michelot 2022; Pohle et al. 2023) or irregular sampling rates (Munden et al. 2021) could eventually be incorporated into the simulator for even greater realism.

## Acknowledgements

We thank Christen H. Fleming for pointing out how to implement home ranging in iSSFs. JF was supported by the National Aeronautics and Space Administration award 80NSSC21K1182 and received partial salary support from the Minnesota Agricultural Experimental Station.

## Authors contributions

JS, TA, BR and US conceived the simulator. US derived the transformation from polar to Euclidean coordinates. All authors contributed to the application of the simulator. JS led the writing with support from all others. JS, BS, and BR implemented and tested the simulator in R.

## Table

Overview of the main functionality of the amt-simulator. Function names are in **bold**. The most important arguments are listed below with their default values in *italics*.

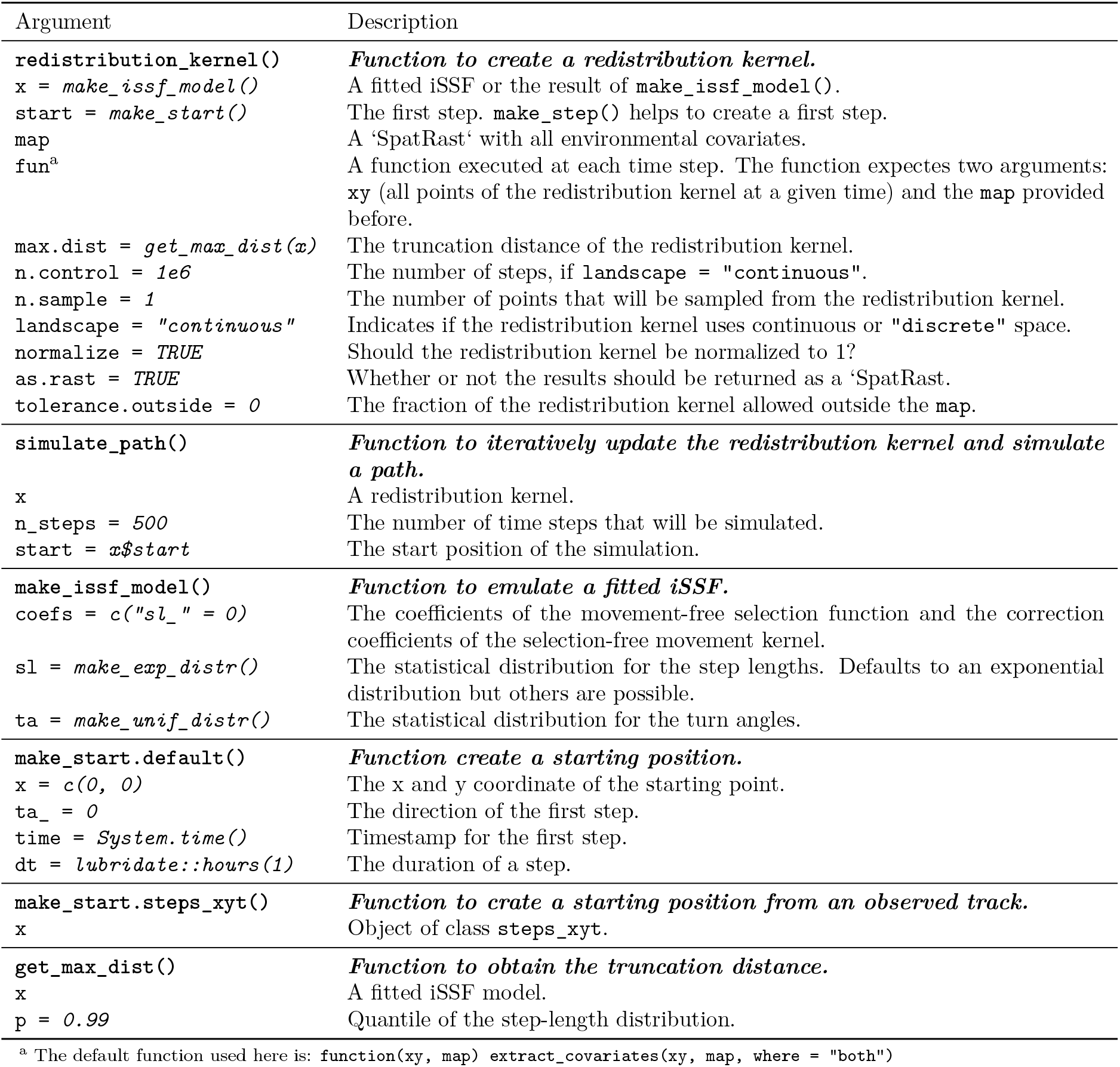

| Argument | Description |
| --- | --- |
| **redistribution_kernel()** | ***Function to create a redistribution kernel.*** |
| `x = make_issf_model()` | A fitted iSSF or the result of `make_issf_model()`. |
| `start = make_start()` | The first step. `make_step()` helps to create a first step. |
| `map` | A 'SpatRast' with all environmental covariates. |
| `fun<sup>a</sup>` | A function executed at each time step. The function expects two arguments: `xy` (all points of the redistribution kernel at a given time) and the `map` provided before. |
| `max.dist = get_max_dist(x)` | The truncation distance of the redistribution kernel. |
| `n.control = 1e6` | The number of steps, if `landscape = "continuous"`. |
| `n.sample = 1` | The number of points that will be sampled from the redistribution kernel. |
| `landscape = "continuous"` | Indicates if the redistribution kernel uses continuous or "discrete" space. |
| `normalize = TRUE` | Should the redistribution kernel be normalized to 1? |
| `as.rast = TRUE` | Whether or not the results should be returned as a 'SpatRast'. |
| `tolerance.outside = 0` | The fraction of the redistribution kernel allowed outside the `map`. |
| **simulate_path()** | ***Function to iteratively update the redistribution kernel and simulate a path.*** |
| `x` | A redistribution kernel. |
| `n_steps = 500` | The number of time steps that will be simulated. |
| `start = x$start` | The start position of the simulation. |
| **make_issf_model()** | ***Function to emulate a fitted iSSF.*** |
| `coefs = c("sl_" = 0)` | The coefficients of the movement-free selection function and the correction coefficients of the selection-free movement kernel. |
| `sl = make_exp_distr()` | The statistical distribution for the step lengths. Defaults to an exponential distribution but others are possible. |
| `ta = make_unif_distr()` | The statistical distribution for the turn angles. |
| **make_start.default()** | ***Function create a starting position.*** |
| `x = c(0, 0)` | The x and y coordinate of the starting point. |
| `ta_ = 0` | The direction of the first step. |
| `time = System.time()` | Timestamp for the first step. |
| `dt = lubridate::hours(1)` | The duration of a step. |
| **make_start.steps_xyt()** | ***Function to crate a starting position from an observed track.*** |
| `x` | Object of class `steps_xyt`. |
| **get_max_dist()** | ***Function to obtain the truncation distance.*** |
| `x` | A fitted iSSF model. |
| `p = 0.99` | Quantile of the step-length distribution. |

<sup>a</sup> The default function used here is: `function(xy, map) extract_covariates(xy, map, where = "both")`

