## Supplement 1 for "Simulating animal space use from fitted integrated Step-Selection Functions (iSSF)"

August 2023

### Transformation from polar to euclidean coordinates

We transform the selection-free movement kernel from a polar coordinate system (where each step is characterized by a step length $\ell$ from a step-length distribution $S$ and bearing $\alpha$, i.e. the absolute angle, from a distribution of bearings $B$) to an Euclidean coordinate system (where each step is characterized by the start $\mathbf{s_1} = (x_1, y_1)$ and end coordinates $\mathbf{s_2} = (x_2, y_2)$). We often use the gamma distribution for $S$ and the von Mises distribution for $B$.

The selection-free movement kernel $\Phi(\ell, \alpha)$ is defined as

$$\Phi(\ell, \alpha) = p_{\text{Gamma}}(\ell) p_{\text{vonMises}}(\alpha)$$

where $p_{\text{Gamma}}(\ell)$ and $p_{\text{vonMises}}(\alpha)$ are the probability density functions of a gamma distribution and von Mises distribution, respectively. To obtain $\Phi(\cdot)$ in Euclidean space, we apply the change-of-variable technique for two random variables. Generally, this states that the joint probability density function for transformed random variables $U = g_1(X, Y)$ and $V = g_2(X, Y)$ is given by $f_{u,v} = f_{x,y}(h_1(u, v), h_2(u, v))|J|$, where $h_1$ and $h_2$ are the inverse transformations for the variables and $|J|$ is the determinant of the Jacobian matrix.

$$\mathbf{J} = \begin{bmatrix} \frac{\partial h_1}{\partial l} & \frac{\partial h_1}{\partial b} \\ \frac{\partial h_2}{\partial l} & \frac{\partial h_2}{\partial b} \end{bmatrix}$$

For our specific case, we have two random variables: the step-length distribution ($\ell$) and bearing ($\alpha$). We apply the following transformation:

- $s_1 = \ell \cos(\alpha) = g_1(\ell, \alpha)$
- $s_2 = \ell \sin(\alpha) = g_2(\ell, \alpha)$

We need the inverse Jacobian matrix for this transformation. We can use the fact that the Jacobian of the inverse transformation is the reciprocal of the orignal transformation.

The Jacobian for this transformation is given by

$$\mathbf{J} = \begin{bmatrix} \cos(\alpha) & -\ell \sin(\alpha) \\ \sin(\alpha) & \ell \cos(\alpha) \end{bmatrix}$$

Next, we take the determinant:

$$
\begin{aligned}
|\mathbf{J}| &= \det(\mathbf{J}) \\
&= \ell\cos(\alpha)^2 + \ell\sin(\alpha)^2 \\
&= \ell\,\underbrace{\left(\cos(\alpha)^2 + \sin(\alpha)^2\right)}_{=1} \\
&= \ell
\end{aligned}
$$

and we use the reciprocal, thus we have $\frac{1}{|\mathbf{J}|} = \frac{1}{S}$ for our case, and the transformation we need to apply is

$$
f_{s_1,s_2}(s_1, s_2) = f_{\ell,\alpha}(\ell, \alpha)\frac{1}{\ell}
$$

Note, that we worked with the bearing here (absolute direction, where East represents zero). For simulating from fitted iSSFs we work with relative turn angles, which can be seen as a rotation of the bearing.
