## Supplementary material for "Simulating animal space use from fitted integrated Step-Selection Functions (iSSF)": Supplment 2: S2_R_code.R

```
#+ message=FALSE
library(lubridate)
library(terra)
library(tidyverse)
library(conflicted)
library(ggplot2)
library(scico)
library(patchwork)
library(amt) # dev version needed: remotes::install_github("jmsigner/amt")
library(here) # here::here() should point to the folder "supplement2" containing 
              # data/cilla.rds and data/env_covar.rds
library(tictoc)


conflict_prefer("select", "dplyr")
conflict_prefer("filter", "dplyr")
dir.create(here::here("ms/figs"), recursive = TRUE, showWarnings = FALSE)

# Simulation case study --------------------------------------------------------

r <- rast(xmin = -10, xmax = 10, ymin = -10, ymax = 10, res = 0.01)
r[] <- 0
p <- terra::vect(cbind(c(1, 1, 6, 6, 1), c(-4, 4, 4, -4, -4)), type = "polygon")
r <- rasterize(p, r, background = 0)
plot(r)
names(r) <- "x"

x <- seq(0.1, 10, 0.1)
plot(x, dgamma(x, shape = 2, scale = 2), type = "l")


set.seed(123)
start <- make_start(c(0, 0), start = 0)
m <- make_issf_model(sl = make_gamma_distr(shape = 2, scale = 2))
rdk1 <- redistribution_kernel(m, start = start, map = r, as.rast = TRUE, 
                              max.dist = 5, landscape = "discrete", 
                              compensate.movement = TRUE)
terra::plot(rdk1$redistribution.kernel)

m <- make_issf_model(sl = make_gamma_distr(shape = 2, scale = 2), 
                     ta = make_vonmises_distr(kappa = 4))
rdk2 <- redistribution_kernel(m, start = start, map = r, as.rast = TRUE, 
                              max.dist = 5, landscape = "discrete", 
                              compensate.movement = TRUE)
plot(rdk2$redistribution.kernel)

start <- make_start(c(0, 0), ta_ = pi/4)
m <- make_issf_model(sl = make_gamma_distr(shape = 2, scale = 2), 
                     ta = make_vonmises_distr(kappa = 4))
rdk3 <- redistribution_kernel(m, start = start, map = r, as.rast = TRUE, 
                              max.dist = 5, landscape = "discrete", 
                              compensate.movement = TRUE)
plot(rdk3$redistribution.kernel)

start <- make_start(c(0, 0), ta_ = 0)
m <- make_issf_model(coefs = c(x_end = 1), sl = make_gamma_distr(shape = 2, 
                                                                 scale = 2), 
                     ta = make_vonmises_distr(kappa = 5))
rdk4 <- redistribution_kernel(m, start = start, map = r, as.rast = TRUE, 
                              max.dist = 5, landscape = "discrete", 
                              compensate.movement = TRUE)
plot(rdk4$redistribution.kernel)

start <- make_start(c(0, 0), ta_ = 0)
m <- make_issf_model(coefs = c(x_end = -1), sl = make_gamma_distr(shape = 2, 
                                                                  scale = 2), 
                     ta = make_vonmises_distr(kappa = 5))
rdk5 <- redistribution_kernel(m, start = start, map = r, as.rast = TRUE, 
                              max.dist = 5, landscape = "discrete", 
                              compensate.movement = TRUE)
plot(rdk5$redistribution.kernel)

terra::plot(r)
points(0, 0)
start <- make_start(c(0, 0), ta_ = 0)
m <- make_issf_model(coefs = c(x_end = 0, "cos(ta_):x_start" = -4), 
                     sl = make_gamma_distr(shape = 2, scale = 2), 
                     ta = make_vonmises_distr(kappa = 4))
rdk6 <- redistribution_kernel(m, start = start, map = r, as.rast = TRUE, 
                              max.dist = 5, landscape = "discrete", 
                              compensate.movement = TRUE)
plot(rdk6$redistribution.kernel)

terra::plot(r)
points(2, 0)
start <- make_start(c(2, 0), ta_ = 0)
m <- make_issf_model(coefs = c(x_end = 0, "cos(ta_):x_start" = -4), 
                     sl = make_gamma_distr(shape = 2, scale = 2), 
                     ta = make_vonmises_distr(kappa = 4))
rdk7 <- redistribution_kernel(m, start = start, map = r, as.rast = TRUE, 
                              max.dist = 5, landscape = "discrete", 
                              compensate.movement = TRUE)
plot(rdk7$redistribution.kernel)

plot_rdk <- function(k, x.lab = NULL, y.lab = NULL, 
                     p = data.frame(x = c(1, 1, 6, 6, 1), 
                                    y = c(-4, 4, 4, -4, -4))) {
  xy <- as.data.frame(k$redistribution.kernel, xy = TRUE)
  pl <- ggplot(xy, aes(x, y, fill = w)) + geom_raster() 
    if (!is.null(p)) pl <- pl + geom_path(aes(x, y), data = p, 
                                          inherit.aes = FALSE, lty = 2, 
                                          col = "grey70") 
    pl + coord_equal() + 
    theme_light() +
    theme(legend.position = "none") +
    scale_fill_scico(palette = "lajolla") +
    labs(x = x.lab, y = y.lab)
}

p1 <- plot_rdk(rdk1, p = NULL, y.lab = "y") + scale_x_continuous(
  limits = c(-5, 6)) + 
  theme(axis.title.x = element_blank(), axis.text.x = element_blank()) +
  labs(subtitle = bquote(b[1]~"= 0; "~b[4]~"= 0; "~b[5]~"= 0"))

p2 <- plot_rdk(rdk2, p = NULL) + scale_x_continuous(limits = c(-5, 6)) + 
  theme(axis.title = element_blank(), axis.text = element_blank()) +
  labs(subtitle = bquote(
    b[1]~"= 0; "~b[4]~"= 4; "~b[5]~"= 0"))
    
p3 <- plot_rdk(rdk4) + scale_x_continuous(limits = c(-5, 6)) +
  theme(axis.title = element_blank(), axis.text = element_blank()) +
  labs(subtitle = bquote(b[1]~"= 1; "~b[4]~"= 4; "~b[5]~"= 0"))

p4 <- plot_rdk(rdk5, y.lab = "y", x.lab = "x") + scale_x_continuous(
  limits = c(-5, 6)) +
  labs(subtitle = bquote(b[1]~"= -1; "~b[4]~"= 4; "~b[5]~"= 0"))

p5 <- plot_rdk(rdk6, x.lab = "x") + scale_x_continuous(limits = c(-5, 6)) +
  theme(axis.title.y = element_blank(), axis.text.y = element_blank()) +
  labs(subtitle = bquote(b[1]~"= 0; "~b[4]~"= 4; "~b[5]~"= -4"))

p6 <- plot_rdk(rdk7, x.lab = "x") + scale_x_continuous(limits = c(-5, 6)) +
  theme(axis.title.y = element_blank(), axis.text.y = element_blank()) +
  labs(subtitle = bquote(b[1]~"= 0; "~b[4]~"= 4; "~b[5]~"= -4"))

(p1 + p2 + p3) / (p4 + p5 + p6) + patchwork::plot_annotation(tag_levels = "a", 
                                                             tag_suffix = ")")

ggsave(here::here("ms/figs/fig1_redistribution_kernels.png"), width = 25, 
       height = 18, units = "cm")


# Figure 2 ----------------------------------------------------------------

set.seed(123)
r <- rast(xmin = -100, xmax = 100, ymin = -100, ymax = 100, res = 0.5)
r
r[] <- 0

p <- terra::vect(cbind(c(1, 1, 50, 50, 1), c(-40, 40, 40, -40, -40)), 
                 type = "polygon")
r <- rasterize(p, r, background = 0)
plot(r)
names(r) <- "x"

start <- make_start(c(0, 0), ta_ = 0)
m <- make_issf_model(coefs = c(x_end = 2), sl = make_gamma_distr(shape = 2, 
                                                                 scale = 2), 
                     ta = make_vonmises_distr(kappa = 1))
rdk.1a <- redistribution_kernel(m, start = start, map = r, landscape = "continuous", 
                                max.dist = 5, n.control = 1e4)
simulate_path(rdk.1a, n.steps = 20)

n <- 50
system.time(p1 <- replicate(n, simulate_path(rdk.1a, n = 30), simplify = FALSE))

# UD
uds <- lapply(c(5, 30), function(i) {
  tibble(
    rep = 1:n, 
    path = map(p1, ~ dplyr::slice(.x, i))
  ) |> unnest(cols = path) |> filter(!is.na(x_)) |> 
    make_track(x_, y_) |> hr_kde(trast = r)
})

pl1 <- tibble(
  rep = 1:n, 
  path = p1
) |> unnest(cols = path) |> 
  ggplot(aes(x_, y_, group = rep)) + geom_path(alpha = 0.1) +
  geom_path(aes(x, y), data = geom(p, df = TRUE), lty = 2, col = "grey30", 
            inherit.aes = FALSE)  +
  geom_point(x = 0, y = 0, col = "red") +
  theme_light() +
  coord_equal() +
  scale_y_continuous(limits = c(-80, 80)) + scale_x_continuous(
    limits = c(-80, 80)) + labs(y = "y", x = "x")

pl2 <- tibble(
  rep = 1:n, 
  path = map(p1, ~ dplyr::slice(.x, 30))
) |> unnest(cols = path) |> 
  ggplot(aes(x_, y_, group = rep)) + geom_point(alpha = 1/10) +
  geom_path(aes(x, y), data = geom(p, df = TRUE), lty = 2, col = "grey30", 
            inherit.aes = FALSE)  + 
  geom_point(x = 0, y = 0, col = "red") +
  theme_light() +
  coord_equal() +
  labs(x = "x") +
  scale_y_continuous(limits = c(-80, 80)) + scale_x_continuous(
    limits = c(-80, 80)) + 
  theme(axis.title.y = element_blank(), axis.text.y = element_blank())

    
xy <- as.data.frame(uds[[2]]$ud, xy = TRUE)
names(xy) <- c("x", "y", "w")
pl3 <- ggplot(xy, aes(x, y, fill = w)) + geom_raster()  +
  geom_path(aes(x, y), data = geom(p, df = TRUE), lty = 2, col = "grey30", 
            inherit.aes = FALSE) +
  coord_equal() + 
  theme_light() +
  geom_point(x = 0, y = 0, col = "red") +
  theme(legend.position = "none") +
  scale_fill_scico(palette = "lajolla") +
  scale_y_continuous(limits = c(-80, 80)) + scale_x_continuous(
    limits = c(-80, 80)) + 
  theme(axis.title.y = element_blank(), axis.text.y = element_blank()) +
  labs(x = "x")

    
pl1 + pl2 + pl3 + plot_annotation(tag_levels = "a", tag_suffix = ")")

ggsave(here::here("ms/figs/fig3_paths.png"), width = 25, height = 10, units = "cm")


# Buffalo study ----------------------------------------------------------------
set.seed(123333)

cilla <- read_rds(here::here("data/cilla.rds"))
env <- rast(read_rds(here::here("data/env_covar.rds")))
env

# Create random steps
ssf_cilla <- cilla %>% steps_by_burst() %>% random_steps(n_control = 100) %>% 
  extract_covariates(env, where = "both") %>% 
  mutate(hour = hour(t1_) + minute(t1_) / 60) %>% 
  filter(complete.cases(.))

ssf_cilla <- cilla %>% steps_by_burst() %>% 
  random_steps(n_control = 100) |> 
  extract_covariates(env, where = "both") %>% 
  mutate(hour = hour(t1_) + minute(t1_) / 60) %>% 
  filter(complete.cases(.))

m_0 <- fit_clogit(ssf_cilla, case_ ~ cos(ta_) + sl_ + log(sl_) + 
                    water_dist_end + strata(step_id_))

# Make check if start is on the map
# This should fail
# redistribution_kernel(m_0, map = terra::rast(env), landscape = "continuous", 
# tolerance.outside = 0.1)

summary(m_0)

start <- make_start(ssf_cilla[1, ])
k1 <- redistribution_kernel(m_0, map = env, start = start)
k1 <- redistribution_kernel(m_0, map = env, start = start,
                      landscape = "continuous", tolerance.outside = 0.2, 
                      n.control = 1e4)

tictoc::tic()
set.seed(1453)
s1 <- simulate_path(k1, n.steps = 1000)
tictoc::toc()

terra::plot(env[["elev"]])
lines(cilla$x_, cilla$y_)
lines(s1$x_, s1$y_, col = "red")


# Home ranging behavior (thanks to Chris Flemming)
m_1 <- fit_clogit(ssf_cilla, case_ ~ cos(ta_) + sl_ + log(sl_) + 
                    water_dist_end + x2_ + y2_ + I(x2_^2 + y2_^2) + 
                    strata(step_id_))
summary(m_1)

k2 <- redistribution_kernel(m_1, map = env, start = make_start(ssf_cilla[1, ]),
                            landscape = "continuous", tolerance.outside = 0.01, 
                      n.control = 1e4)
set.seed(18)
s2 <- simulate_path(k2, n.steps = 1000)

terra::plot(env[["elev"]])
lines(cilla$x_, cilla$y_)
lines(s2$x_, s2$y_, col = "red")


## Barriers
water <- env[["water_dist"]] > 100
water <- crop(water, ext(water) - 5000)
plot(water)
ww <- patches(water, zeroAsNA = TRUE)
plot(ww)
ww[ww == 3] <- 2
names(ww) <- "water_crossed"
env1 <- c(ww, crop(env, ww))

ssf_cilla <- cilla %>% steps_by_burst() %>% random_steps(n_control = 100) %>% 
  extract_covariates(env1, where = "both") %>% 
  mutate(hour = hour(t1_) + minute(t1_) / 60) %>% 
  filter(complete.cases(.))

m_3 <- fit_clogit(ssf_cilla, case_ ~ cos(ta_) + sl_ + log(sl_) + 
                    water_dist_end + x2_ + y2_ + I(x2_^2 + y2_^2) +
                    I(water_crossed_end != water_crossed_start) + 
                    strata(step_id_))
summary(m_3)

set.seed(18)
k3 <- redistribution_kernel(m_3, map = env1, start = make_start(ssf_cilla[1, ]),
                            landscape = "continuous", tolerance.outside = 0.01, 
                      n.sample = 100, n.control = 1e4)

s3 <- simulate_path(k3, n.steps = 1000)

terra::plot(env1[["water_crossed"]])
lines(cilla$x_, cilla$y_)
lines(s3$x_, s3$y_, col = "red")

rast <- terra::as.data.frame(env1[["water_crossed"]], xy = TRUE)

# Figure 3 ------------------
x.mean <- cilla$x_ |> mean()
y.mean <- cilla$y_ |> mean()

cilla1 <- cilla
cilla1$x_ <- (cilla1$x_ - x.mean) / 1e3
cilla1$y_ <- (cilla1$y_ - y.mean) / 1e3

rast$x <- (rast$x - x.mean) / 1e3
rast$y <- (rast$y - y.mean) / 1e3

bind_rows(
  s1 |> mutate(model = "Model 1"),
  s2 |> mutate(model = "Model 2"),
  s3 |> mutate(model = "Model 3")
) |> 
  mutate(x_ = (x_ - x.mean) / 1e3, 
         y_ = (y_ - y.mean) / 1e3) |> 
  ggplot(aes(x_, y_)) + 
  geom_raster(aes(x, y, fill = factor(water_crossed)), data = rast, 
              inherit.aes = FALSE) +
  geom_path(data = cilla1, col = "white", alpha = 0.6) +
  geom_path(col = "red") +
  facet_wrap(~ model) +
  coord_equal() + 
  theme_light() +
  theme(legend.position = "none") +
  scale_fill_manual(values = c("grey30", "grey70")) +
  lims(x = c(-15, 25), y = c(-22, 15)) +
  labs(x = "x", y = "y")

ggsave(here::here("ms/figs/african_buffalo1.png"), width = 25, height = 10, 
       units = "cm")

sessionInfo()
```
